# Retrieval-extinction within the memory reconsolidation window does not influence appetitive choice

**DOI:** 10.1101/014316

**Authors:** Akram Bakkour, Tom Schonberg, Ashleigh M. Hover, Russell A. Poldrack

## Abstract

Influencing choice behavior is key to achieving behavioral change. Traditional means to influence choice behavior rely on effortful self control, which is known to be fragile under several circumstances, rendering these methods ineffectual in maintaining any change in behavior over time. Behavioral maintenance efforts are likely more effective over the long term if they target more automatic processes such as attention or memory. Memories are not set in stone and are vulnerable to change and updating under certain circumstances when retrieved. It is possible to target specific memories for updating. In two studies, we sought to update the memory for an appetitive choice by way of reversal learning following retrieval of the targeted choice behavior. We found that targeting memories of a choice behavior for updating shortly after a reminder did not significantly attenuate the renewal of the targeted choice under extinction conditions. Possible explanations and suggested future directions are discussed.

## Introduction

When retrieved, established memories enter a labile state that renders them susceptible to modification, providing an opportunity to strengthen, weaken or update memories. These memories then re-stabilize through a process known as reconsolidation in order to persist (Nader et al., 2000). Reconsolidation has been shown to be protein-sythesis dependent and transient. Amnestic agents such as anisomycin applied at least ten minutes, but no more than six hours after memory retrieval blocked the return of fear (Duvarci and Nader, 2004). Memory updating during reconsolidation has been demonstrated in many species using several different memory paradigms, suggesting that this process is a fundamental feature that spans different kinds of memory (see Besnard et al., 2012; Alberini and Ledoux, 2013; Reichelt and Lee, 2013, for review).

Important potential clinical applications for memory updating during reconsolidation have been proposed, for example in the treatment of post-traumatic stress disorder (PTSD, Debiec and Ledoux, 2006; Brunet et al., 2008). Patients with PTSD who were given the beta-adrenergic blocker propranolol shortly after being asked to describe the traumatic event they had experienced showed a marked decrease in physiological responding during traumatic script-driven imagery one week later (Brunet et al., 2011). Non-invasive behavioral retrieval-extinction of fear within the reconsolidation window has also been shown to be effective in the return of fear following Pavlovian cued fear conditioning (Monfils et al., 2009; Schiller et al., 2010).

Although there have been several successful efforts to interfere with memories during reconsolidation in order to prevent the return of fear, relatively few have focused on updating appetitive behavior. These efforts have mostly focused on appetitive Pavlovian conditioning such as drug use (Milton et al., 2008; Lee and Everitt, 2008b). Administration of propranolol had a limited effect on the treatment of drug addiction (Milton et al., 2012). However, retrieval-extinction manipulations have proven useful for reducing conditioned place preference for morphine and cocaine in rats as well as cue-induced heroin craving in humans (Xue et al., 2012). Behavioral interference with a memory within the reconsolidation window could prove useful for updating maladaptive behavior in favor of improved behavior. Many of the successful efforts to update maladaptive memories however have targeted Pavlovian rather than instrumental memories.

Exton-McGuinness et al. (2014) have shown that well-learned instrumental memories can be disrupted by administration of a noncompetitive N-methyl-D-aspartate receptor (NMDAR) antagonist in rats. Previous studies have failed to demonstrate disruption of instrumental memories during reconsolidation (Hernandez and Kelley, 2004; Mierzejewski et al., 2009), but the discrepancies in the findings are likely due to differences in the parameters during the reactivation session necessary to destabilize the memory (Piñeyro et al., 2014). Censor et al. (2014) recently showed that procedural memory in humans is susceptible to updating when applying repetitive transcranial magnetic stimulation (rTMS). Fewer studies have employed a behavioral retrieval-extinction paradigm during reconsolidation to target procedural memories for updating.

Walker et al. (2003) trained human participants on a motor sequence task and 24 hours later trained the participants on a new motor sequence immediately after brief rehearsal of the sequence from the previous day. They found that speed and accuracy for the day 1 sequence decreased as a result of the interference of the second motor sequence training when the latter was performed immediately, but not when performed six hours after the reminder. However, to our knowledge, no study has targeted memories for contingencies that govern choice behavior for updating using a behavioral retrieval-extinction paradigm.

In this study, we employ an ABA renewal paradigm where training is first conducted in context A, reversal learning is conducted in context B, then an extinction test is conducted in context A to test whether reversal learning within the reconsolidation window reduces renewal of the first-learned response. This approach has clear implications for lasting behavioral change.

## Materials and Methods

### Participants

70 healthy participants completed two studies. Participants were placed into one of two experimental conditions. They started reversal training on day 2 either 10 min or 6 hr after a reminder of day 1 training stimuli (for details please see Task section below). Table 1 summarizes participant demographic details for the two studies. Sample sizes are similar to previously published studies.

**Table 1.**
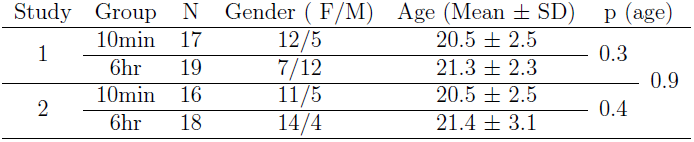
Demographic details for studies 1 & 2. P values reflect signifícance in two-sided independent samples t-tests for age. SD (Standard Deviation).

All participants were right handed, had normal or corrected-to-normal vision, no history of psychiatric or neurologic disease and were not taking any medication that would interfere with the experiment. Participants agreed to participate in the study over three consecutive days and gave informed consent. The study was approved by the institutional review board (IRB) at the University of Texas at Austin.

### Task

Participants agreed to come in for four study sessions at two locations over three consecutive days in two studies. Figure 1 outlines the design for studies 1 and 2. All details were identical for both studies, except for the number of stimuli and the number of times each stimulus was repeated. These differences are summarized in Table 2.

**Figure 1.**
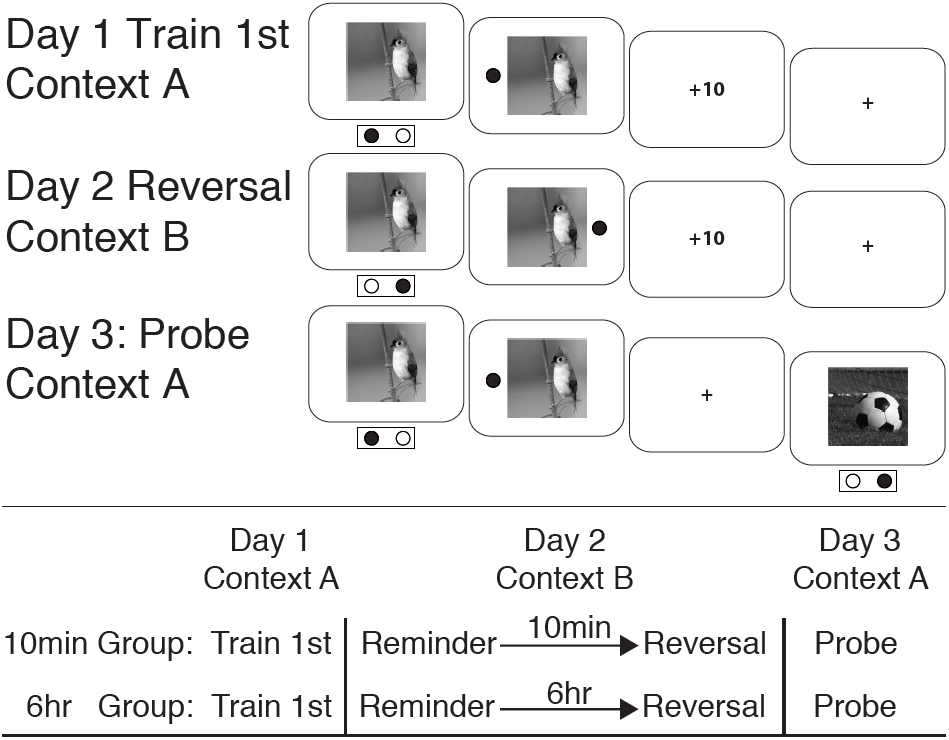
Schematic of experimental procedure for memory interference studies. Details of each step are described in the Task section. Briefly, participants learn to press one of two buttons to earn points 80% vs. 20% of the time on day 1 in context A (room A in building A). On day 2, contingencies are reversed in a new context B (room B in building B) either ten minutes or six hours after a reminder. On day 3, participants perform a probe under extinction conditions in context A.

**Table 2.** Differences in the number of stimuli and the number of times each stimulus was presented during the day 1 and day 2 training phases of both studies.

| Study | # of stimuli | # of times each stimulus was presented on day 1 | # of times each stimulus was presented on day 2 |
| --- | --- | --- | --- |
| 1 | 8 | 10 | 5 |
| 2 | 4 | 8 | 4 |

### Train day 1

On the first visit, participants learned to press one of two buttons to earn points 80% rather than 20% of the time in context A. This was in room A in building A and the computer screen background color was A. Half the stimuli were associated with a more favorable right button press and the other half with a left button press. Stimuli were black and white photographs of neutral objects. The number of stimuli and the number of times each stimulus was repeated are summarized in Table 2. Stimuli appeared in random order per block of eight or four presentations so that the time between presentations of a particular stimulus remained relatively constant.

### Reversal day 2

On the second visit, participants went to a different room in a different building (context B) and filled out a computer adapted version of BIS-11 (Patton et al., 1995) then were reminded of the stimuli with a single exposure to each of the stimuli. To ensure that the participants were attending to the stimuli, they were asked to press a different button (the space bar) when a stimulus appeared on the screen. Participants were assigned to one of two groups. One group received the reminder six hours before reversal training on the second day and the other ten minutes before reversal training on the second day. These times were shown to correspond to times outside and within the reconsolidation window respectively (Duvarci and Nader, 2004; Monfils et al., 2009). During training on the second day, the first day contingencies were switched and participants learned to reverse what they had learned the previous day. Each stimulus appeared five times for study 1 and four times for study 2 during reversal training on day 2. We chose to reduce the training by half from day 1 during reversal training on day 2 to avoid overtraining effects during reversal since participants learn faster during reversal learning compared to initial leaning (pilot data not shown). We aimed for learning on day 1 and day 2 to be equivalent.

### Probe day 3

On the third day, participants returned to room A in building A that they had visited on day 1 (context A). They were presented with the same items as the previous two days and asked to press one of two buttons, but no outcome was provided. Participants were told that although they would receive no outcome information, the computer would continue to count points in the background and that it was important for them to press the button they thought would yield points. Renewal was measured as choices consistent with first-learned contingencies on day 1. This ABA renewal paradigm was designed to detect updating of contingency memory trace when training after a contingency switch occurs within the reconsolidation window (i.e. ten minutes after a reminder).

### Analysis

#### Training

To test for successful training to press one button over another for each stimulus, we ran repeated measures logistic regression on the odds of choosing the high-reward response (that yields points 80% of the time) to the low-reward response (that yields points 20% of the time) on the last block vs. the first block of presentations during training on day 1 with a grouping factor for participant. We ran the same regression model for reversal training on day 2. We also ran regression to compare the groups (10min/6hr) on the last block of presentations on each day to test for any differences in learning between the groups.

#### Probe

We performed repeated measures logistic regression on the odds of choosing the high-reward response from day 1 to the high-reward response from day 2 against equal odds with a participant grouping factor separately for the 10min and 6hr groups. These regressions test for renewal of the first-learned behavior (high-reward response on day 1) during probe on day 3 (which takes part in context A, the same as training on day 1). We also ran repeated measures logistic regression to test for the difference between the 10min and 6hr groups on the odds of performing the first-learned to second-learned behavior. We hypothesized that the 6hr group (i.e. those who received reversal training outside the reconsolidation window) would exhibit significant renewal compared to the 10min group (i.e. those whose reversal training occurred within the reconsolidation window and whose memory for first-learned contingencies were targeted for updating by contingencies on day 2).

The influence of successful training on day 1 on renewal and its interaction with group assignment were tested using linear regression. We hypothesized that the influence of training from day 1 on renewal would be weaker for the 10min group compared to the 6hr group, which would suggest that the memory for contingencies from day 1 was updated by contingencies from day 2 in the 10min group.

## Results

### Training

Participants learned to choose the high-reward option (button press associated with 80% reward) by the end of training on day 1 and learned to switch their responses on day 2 (Figure 2A and C and Table 3). There were no differences in choice of high-reward option between the 6hr and 10min groups on the last block of presentations on either day or in either study, suggesting that learning was equivalent between the groups on both days. There were also no differences in reaction time during training between the groups (p’s > 0.15).

**Figure 2.**
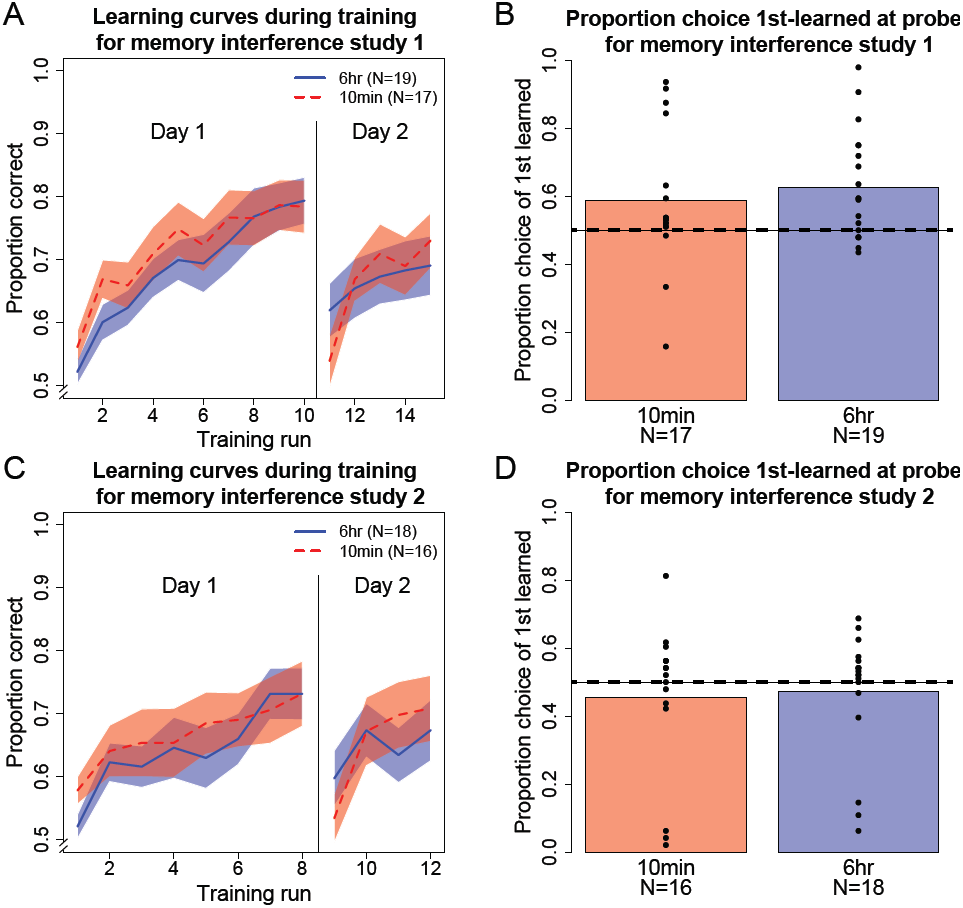
Behavioral results for memory interference studies. The top row (A,B) shows resuits from study 1. Participants were trained on eight stimuli (N = 36). The bottom row (C,D) shows results from study 2. Participants were trained on four stimuli (N = 34). A,C) The solid blue line represents the mean proportion of correct responses (deSned as a high-reward response yielding points 80% of the time) over runs of the task for the 6hr group. The interrupted red line shows the same for the 10min group (group Ns in legend on top right). The shaded area represents one standard error of the mean (SEM). B,D) The bars represent the mean proportion of first-learned response (high-reward response from day 1) during the probe phase under extinction conditions. The left red bar includes participants in the 10min group, the right blue bar includes participants in the 6hr group. The dots represent the proportion of first-learned responses for each individual participant.

**Table 3.**
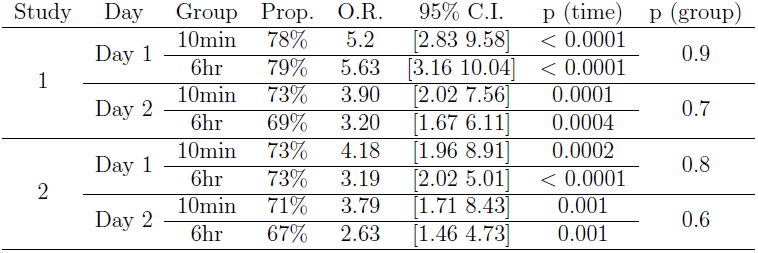
Descriptive statistics for training phase behavior in studies 1 & 2. Proportion (Prop.) choice of high-reward option at the end of training on each day. Odds ratio (O.R) for choice of high-to low-reward option at the end of training on each day. Confidence interval (C.I) on odds ratio and p-value (time) for odds of choosing high-reward option at the end of training vs. the beginning of training and p-value (group) for the odds of choosing high-to low-reward option at the end of training between groups.

| Study | Day | Group | Prop. | O.R. | 95% C.I. | p (time) | p (group) |
| --- | --- | --- | --- | --- | --- | --- | --- |
| 1 | Day 1 | 10min | 78% | 5.2 | [2.83 9.58] | < 0.0001 | 0.9 |
|  |  | 6hr | 79% | 5.63 | [3.16 10.04] | < 0.0001 |  |
|  | Day 2 | 10min | 73% | 3.90 | [2.02 7.56] | 0.0001 | 0.7 |
|  |  | 6hr | 69% | 3.20 | [1.67 6.11] | 0.0004 |  |
| 2 | Day 1 | 10min | 73% | 4.18 | [1.96 8.91] | 0.0002 | 0.8 |
|  |  | 6hr | 73% | 3.19 | [2.02 5.01] | < 0.0001 |  |
|  | Day 2 | 10min | 71% | 3.79 | [1.71 8.43] | 0.001 | 0.6 |
|  |  | 6hr | 67% | 2.63 | [1.46 4.73] | 0.001 |  |

### Probe

Using eight stimuli (study 1), participants in the 6hr group displayed significant renewal of the first-learned response (mean proportion choice of first-learned = 63%, odds of first-to second-learned response p = 0.002). However, renewal of first-learned behavior was not mitigated by reversal learning within the reconsolidation window (mean proportion choice of first-learned = 59%, odds of choosing first-to second-learned in 10min group p = 0.074, and p = 0.57 for the comparison of 6hr to 10min group, Figure 2B). Simplifying the task and using four stimuli did not achieve the hypothesized reduced renewal effect in the 10min group in a second sample of N = 34 (study 2, mean proportion choice of first-learned = 45% vs. 47% for 6hr group, odds of choosing first- over second-learned for 10min vs. 6hr group p = 0.7, Figure 2D). There were no differences in reaction time between the groups at probe (p’s > 0.9).

We also tested the effect of learning on day 1 on renewal of first-learned behavior at probe on day 3. There was a main effect of correct choices at the end of day 1 on renewal of responses at probe (p = 0.0008 for study 1 and 0.05 for study 2), no main effect of group assignment (10min/6hr, p’s > 0.5) and no interaction between responses at the end of day 1 and group assignment on choice at probe (p’s > 0.8), suggesting that there was not significant interference with memory for contingencies from day 1 by conducting reversal learning during the reconsolidation window.

## Discussion

Targeting specific memories for updating during reconsolidation is a promising avenue for treating disorders such as PTSD and drug abuse. In the studies described here, our goal was to interfere with appetitive memory for choice contingencies using reversal learning within the reconsolidation window. We employed an ABA renewal paradigm to test renewal of first-learned responses under extinction conditions after conducting reversal learning either within the reconsolidation window (ten minutes after a reminder of the stimuli) or outside the reconsolidation window (six hours after the reminder). We found that although there is some evidence of significant renewal of first-learned responses when reversal learning was conducted outside the reconsolidation window, there was no evidence of significant updating of memory for first-learned responses or attenuation of renewal of first-learned responses following reversal learning within the reconsolidation window.

Piñeyro et al. (2014) showed that reactivation duration of a fear memory is a critical factor for trace destabilization. More specifically, they found that four minutes, but not one minute of reactivation before the extinction of a fear memory prevented the return of fear. In our studies, reactivation consisted of a single one second presentation of each of the stimuli, which amounted to a maximum of 32 seconds of reactivation. The short reactivation task might not have been sufficient to initiate synaptic protein degradation, which typically takes place at least three minutes after retrieval and is crucial for the destabilization of retrieved memories (Lee et al., 2008).

In our studies, the reminder also involved a different response (press space-bar) than on day 1 training (press one of two buttons for a chance to earn points). Lee and Everitt (2008a) trained rats to associate a lever press with a sucrose reward. Rats were then administered the NMDAR antagonist MK-801 on the second day after a reminder of day 1 training. Reactivation of the lever press-sucrose association was achieved through re-exposure that was either contingent or not on a lever press. Reconsolidation of the lever press-sucrose memory was blocked for rats that received the reminder contingently, but not those who received the reminder noncontingently upon a lever press. The authors concluded that stimuli may have to be presented in the same manner as during training in order to render previously acquired memories unstable. Not performing the same task as on day 1 for the reminder in our studies could have contributed to weaker destabilization of the memory for day 1 contingencies and thus a lack of memory updating during reversal learning.

Spatial context has also been suggested to play an important role in the updating of episodic memories. Hupbach et al. (2008) showed that new learning in the same spatial context as original learning is necessary and sufficient for the incorporation of new information into existing episodic memories. In our studies, the reminder and by design the reversal learning were conducted in a different spatial context than learning on day 1. The switch in spatial context during the reminder on day 2 may have contributed to a lack of full reinstatement and destabilization of the memory for day 1 contingencies in our studies. However, there is evidence that reconsolidation is not triggered when no new information is learned during the reminder trial (Sevenster et al., 2012). Thus it is possible that reminder trials that are identical to the previous day training trials might not provide any new information. In our studies, performing a different response during the reminder trials than on the previous day’s training trials might not provide any new information in the form of a prediction error to trigger reconsolidation. The lack of prediction errors during reminder in our studies might also be a factor in the lack of updating of day 1 contingency memory traces.

Further studies that vary the retrieval task, timing and spatial context during reminder should be conducted to determine the retrieval parameters necessary to destabilize memory for contingencies in the present task, and render it amenable to updating by new contingencies. Although recent years have seen an exponentially growing body of evidence demonstrating reconsolidation of human memories across memory domains (Besnard et al., 2012), not all of the precise boundary conditions that govern reconsolidation across types of memory have been identified and characterized. In a recent review, Schwabe et al. (2014) pose a number of questions for the scientific community to answer relating to human memory reconsolidation. These questions pertain to potential individual differences in the susceptibility to memory updating after retrieval and factors that might influence such differences, the duration of the reconsolidation window and the time over which any memory modifications might last, and the brain mechanisms that support reconsolidation of various types of memory. Although many questions remain to be answered, a better understanding of memory reconsolidation processes has high potential for the treatment of certain psychological and behavioral disorders.

Conict of interest: None of the authors have conicts of interest to report.

## Acknowledgements

This research was generously funded by National Institutes of Health grant 1R01AG041653. The authors would also like to thank Dr. Marie Monfils for helpful suggestions during task design.

